# Connectivity profile laterality in corticostriatal functional circuitry: a fingerprinting approach

**DOI:** 10.1101/2021.06.04.447001

**Authors:** Cole Korponay, Elliot A Stein, Thomas J Ross

## Abstract

An abnormal magnitude of hemispheric difference (i.e. laterality) in corticostriatal circuits is a shared feature of numerous neurodevelopmental and psychiatric disorders. Detailed quantitation and regional localization of corticostriatal laterality in normative samples stands to further the understanding of hemispheric differences in healthy and disease states. Here, we used a fingerprinting approach to quantify “functional connectivity profile laterality” – the overall magnitude by which a voxel’s profile of connectivity with homotopic regions of the ipsilateral and contralateral cortex differs – in the striatum. Laterality magnitude heatmaps revealed “laterality hotspots” – constituting outliers in the voxel-wise distribution –in the right ventrolateral putamen and left central caudate. Findings were replicated in an independent sample, with significant (p<0.05) spatial overlap observed between the location of the laterality hotspots across samples, as measured via Dice coefficients. At both hotspots, a primary driver of overall laterality was the difference in striatal connectivity strength with the right and left pars opercularis of the inferior frontal gyrus. Right and left striatum laterality magnitude maps were found to significantly differ (p<0.05) at the hotspot locations. Moreover, using subjects’ left, but not right, striatum laterality magnitude maps, a support vector machine trained on a discovery sample (n=77) and tested on a replication sample (n=77) significantly predicted (r=0.25, p=0.028) subject performance on a language task, known for its lateralized nature. Laterality magnitude maps remained consistent across different cortical atlas parcellations and did not differ significantly between right-handed and left-handed individuals. In sum, meaningful variation in functional connectivity profile laterality – both spatially within the striatum and across subjects – is evident in corticostriatal circuits. Findings provide a basis to examine corticostriatal connectivity profile laterality in psychiatric illness.

## INTRODUCTION

Asymmetries between the left and right hemispheres of the brain constitute a fundamental feature of neural organization. These asymmetries, which manifest both in gross anatomy and in structural and functional connections, contribute to the neural basis of lateralized brain functions[1], including notably language[2], inhibitory control[3, 4], and salience detection[5]. An abnormal magnitude of hemispheric asymmetry (i.e. laterality) in neural substrates and functional connectivity (FC) is a shared feature of numerous neurodevelopmental and psychiatric disorders, including autism spectrum disorders (ASDs), attention-deficit/hyperactivity disorder (ADHD), and schizophrenia[6, 7]. Individuals with schizophrenia, for instance, have reduced leftward structural[8, 9] and FC [10] asymmetry (i.e., more symmetry) compared to healthy individuals. Common across many of these disorders is atypical asymmetry in frontal cortical-striatal circuitry in particular[6, 11, 12]. Individuals with ASDs have reduced leftward asymmetries or reversed asymmetries in frontal cortical language regions, with more atypical asymmetry associated with greater language deficits[13]. Youths with ADHD have reduced rightward asymmetry in caudate-ventrolateral prefrontal cortex and caudate-dorsolateral prefrontal cortex white matter tract volume compared to controls[11]. In a recent mega-analysis of 22 structural brain MRI datasets, substance dependence was found to be associated with reduced rightward structural asymmetry of the nucleus accumbens[12].

Most prior studies have evaluated laterality by comparing individual pairs of brain regions or connections. A dimension of laterality that has been relatively underexplored is that pertaining to FC profiles. A brain area’s FC profile, or “fingerprint”, is the multivariate set of connection strengths it has with other areas of the brain[14, 15]. The value of examining functional connectivity profiles is that, more so than any one particular connection, it is the combinational makeup of a brain area’s full set of connections that most shapes its activity and function[15]. However, there has been little examination of laterality in FC profiles, including whether its magnitude varies spatially within the brain and whether inter-subject variation relates to capacities in lateralized functions like language. Investigating these questions – particularly in corticostriatal circuitry, where atypical laterality is frequently implicated in neuropsychiatric disease[6, 11, 12] – stands to further the understanding of hemispheric differences in lateralized functions in healthy and disease states.

Anatomical tract-tracing studies in non-human primates indicate that *structural* connectivity profiles in frontal cortical-striatal circuits are highly lateralized[16–18]. Specifically, they find that frontal cortical regions project predominantly to the ipsilateral striatum, and that only a limited number of projections cross the corpus callosum to terminate in the contralateral striatum[16–18]. This suggests that a given striatal area’s structural connectivity profile is heavily weighted toward its ipsilateral connections. Yet, prior functional imaging studies find that the striatum’s *functional* connections with homotopic areas of the ipsilateral and contralateral frontal cortex appear qualitatively comparable[19, 20]. The absence of readily apparent laterality in frontal cortical-striatal *functional* connections likely arises from the strong cortico-cortical connections and correlated activity between homotopic cortical regions, wherein any third region (e.g., the striatum) would be expected to have similar correlated activity with both regions.

Therefore, although absolute levels of FC profile laterality are likely very low throughout the striatum, the possibility remains that there is meaningful variation – both spatially within the striatum and across individuals. Characterization of this variation in a normative sample would provide a novel basis to probe laterality in frontal cortical-striatal circuits in disease states. Furthermore, identification of laterality “hotspots”, where the difference between the ipsilateral and contralateral connectivity profiles is relatively large, may indicate striatal locations that play important roles in lateralized functions. It is also of interest to determine whether left-handedness influences differences in fronto-striatal connectivity profile laterality compared to righthandedness. Given that left-handedness is appreciably more common amongst a number of neurodevelopmental and neuropsychiatry populations that display atypical fronto-striatal FC [8, 13], findings in a normative sample may serve as a basis for understanding the relationship between handedness and abnormal fronto-striatal connectivity in these disease states.

## METHODS

### Overview

Here we propose and implement a modified voxel-wise fingerprinting[14] approach to quantify the laterality magnitude of each striatal voxel’s frontal cortical FC profile. As the foundation for our analytic pipeline, we separately established an ipsilateral and a contralateral frontal cortex connectivity fingerprint for each voxel in the striatum. Each fingerprint encoded the strength of FC between a given striatal voxel and 15 unilateral frontal cortical subregions (‘targets’). The difference between the ipsilateral and contralateral fingerprint at each voxel – measured via Manhattan distance[14] – was operationalized to represent the overall magnitude of laterality in corticostriatal functional connections at each striatal voxel. This allowed for examination of regional variation in laterality within the striatum, and the identification of laterality “hotspots.” We then compared the Manhattan distance (laterality magnitude) at homotopic voxels in the right and left striatum to examine the degree of laterality symmetry. Finally, we examined which frontal cortical subregions were the biggest drivers of connectivity laterality.

We repeated this process in a series of sensitivity and replicability tests. To examine the robustness of the fingerprinting method for assessing laterality magnitude, we examined the effect of using a different cortical atlas parcellation. If the connectivity profile laterality metric is indeed an inherent property of a voxel’s connectivity profile and not an artifact of a specific atlas parcellation, we would expect the voxels identified as having comparatively high laterality to remain largely consistent, regardless of the atlas used. Second, we examined replicability in an independent cohort. Third, we examined differences between right-handed and left-handed individuals. Finally, we assessed whether connectivity profile laterality magnitude was associated with behavioral performance on a task that recruits lateralized functions (i.e., language) and one that does not (i.e., delay discounting).

### Participants

Resting state data used in these analyses are derived from the Human Connectome Project (HCP) Q1-Q6 Data Release. A detailed description of HCP subject recruitment has been provided [21, 22]. Briefly, individuals were excluded by the HCP if they reported a history of major psychiatric disorder, neurological disorder, or medical disorder known to influence brain function. For our discovery sample[23], subjects were also excluded who were related or met the following stringent head motion criteria: (1) range of head motion in any translational direction greater than 1 mm or (2) average scan-to-scan head motion greater than 0.2 mm. After excluding one subject due to artifacts, our final discovery sample consisted of 77 individuals (age: 22-35, 28 males, all right-handed).

### fMRI Data Acquisition

HCP neuroimaging data were acquired with a standard 32-channel head coil on a Siemens 3T Skyra modified to achieve a maximum gradient strength of 100 mT/m[21, 22, 24]. Gradient-echo EPI images were acquired with the following parameters: TR = 720 ms, TE = 33.1 ms, flip angle = 52°, FOV = 280 × 180 mm, Matrix = 140 × 90, Echo spacing = 0.58 ms, BW = 2,290 Hz/Px. Slice thickness was set to 2.0 mm, 72 slices, 2.0 mm isotropic voxels, with a multiband acceleration factor of 8. Resting state data were acquired from two runs of approximately 14.4 min each (REST1). One run was acquired with a right-to-left phase encoding and the other run was acquired with a left-to-right phase encoding. Thus, REST1 is comprised of 28.8 min of data. Participants were instructed to lie still with their eyes open and fixated on a bright crosshair on a dark background.

### Preprocessing

We began with HCP minimally preprocessed resting-state data [25]. Briefly, this preprocessing pipeline removes spatial distortions, realigns volumes to compensate for subject motion, registers the echo planar functional data to the structural data, reduces the bias field, normalizes the 4D image to a global mean, and masks the data with a final FreeSurfer-generated brain mask[25] We further preprocessed these scans including spatial blurring with a 6-mm fullwidth half-maximum Gaussian kernel and temporal filtering (0.01<f <0.1 Hz).

ROIs for the striatal fingerprint were based on the frontal cortical parcellation definitions from the Harvard-Oxford cortical atlas (**Supplemental Figure 1a**), and included: supplementary motor cortex, superior frontal gyrus, subcallosal cortex, precentral gyrus, paracingulate gyrus, middle frontal gyrus, insular cortex, inferior frontal gyrus pars opercularis, inferior frontal gyrus pars triangularis, frontal pole, frontal orbital cortex, frontal operculum cortex, frontal medial cortex, central opercular cortex, and anterior cingulate cortex. Sample-specific right and left striatum masks were created by averaging the union of the caudate, putamen, and nucleus accumbens FreeSurfer parcellations of each subject.

Resting-state functional connectivity (rsFC) was assessed for each frontal cortical seed (ROI) using the mean resting-state BOLD time series, extracted from each participant. The mean time series from each ROI was included in a GLM with 17 additional regressors of no interest: six motion parameters (three translations and three rotations) obtained from the rigid-body alignment of EPI volumes and their six temporal derivatives; the mean time series extracted from white matter; the mean times series extracted from CSF; and a second-order polynomial to model baseline signal and slow drift. To further control for subject motion, volumes were censored for framewise motion displacement (i.e., volume to volume movement) >0.5 mm [26, 27]. The output of R^2^ values from the GLM were converted to Z-scores using the Fisher R-to-Z transform.

### Behavioral Tasks

To study the relationship between connectivity profile laterality and behavior, we examined 1) a task that engages a lateralized functional capacity (e.g. language) and 2) a task that engages a non-lateralized functional capacity (e.g. delay discounting). As such we first examined findings from a reading recognition task in which subjects were asked to read and pronounce letters and words as accurately as possible (oral reading recognition [ReadEng])[28]. Data for the task was obtained in the HCP sample using the NIH toolbox[29]. Second, we examined findings from a delay discounting task[30] that measures the magnitude by which subjects undervalue rewards that are delayed in time. For this task the examined outcome measures were area under the curve for discounting of $200 [DDisc_AUC_200] and area under the curve for discounting of $40,000 [DDisc_AUC_40K][31].

### Analytic Strategy

#### Comparison of ipsilateral and contralateral fingerprints within striatal voxels

In order to compare the ipsilateral and contralateral frontal cortical connectivity fingerprint at each voxel in the striatum, we carried out the following procedure (**Supplemental Figure 2**). First, for each subject, we generated voxel-wise maps of the correlation (Pearson’s *r*) between the mean time series of each frontal cortical ROI and each voxel in the striatum. These Pearson’s *r* maps were transformed to Z-score maps using Fisher’s *r*-to-Z transformation. This resulted in 30 voxel-wise Z-score striatal maps (15 each for the ipsilateral and contralateral hemisphere cortical ROIs). Then, we computed voxel-wise Z-score difference maps by taking the absolute value of the difference between the ipsilateral and contralateral Z-score map of each ROI:

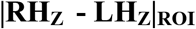

This produced 15 difference maps (one for each ipsilateral - contralateral ROI pair) for each subject. Next, we took the sum of the 15 difference maps, resulting in a final voxel-wise map for each subject whose values represent the Manhattan distance between the ipsilateral and contralateral frontal cortical fingerprint at each striatal voxel:

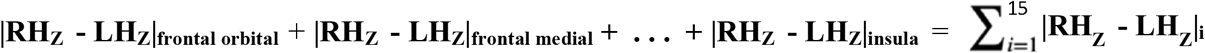

In this way, the Manhattan distance value provides a measure of the magnitude of total frontal cortical FC laterality at each striatal voxel. Finally, we computed the mean of the 77 subject-level Manhattan distance maps, resulting in a group-level voxel-wise map encoding the average within-subject Manhattan distance at each striatal voxel.

Typically, in order to determine a significance threshold for Manhattan distance values derived for fingerprinting analysis, the fingerprints are permuted 10,000 times or more to create a Manhattan distance test statistic distribution[14]. This kind of analysis informs whether any of the empirically measured Manhattan distance values are significantly bigger (or smaller) than would be expected by chance if the data were random. However, this kind of analysis would be uninformative for the purposes of this investigation. As discussed previously, due to the highly correlated activity of homotopic frontal cortical ROIs, any third region (e.g., a striatal voxel) will have a very similar magnitude of connectivity with the LH and RH ROI of a region. As a corollary, then, a striatal voxel’s connectivity with a given region’s LH ROI is likely to be more similar to its connectivity with that same region’s RH ROI than the RH ROI of any other region. The result is that a Manhattan distance calculated by summing the connectivity differences between pairs of matched LH-RH ROIs (e.g., right OFC - left OFC), as opposed to mismatched LH-RH ROIs (e.g., right OFC - left dlPFC), will almost always yield the lowest or close to the lowest possible Manhattan distance value. Since the permutation testing process involves the calculation of Manhattan distances from randomly permuted LH-RH ROI pairs, *a priori* we know that voxels’ empirically measured Manhattan distances will be significantly smaller than nearly all possible values produced from random permutation. As a result, the permutation approach would simply re-confirm that, overall, laterality in frontal cortical-striatal circuits is substantially lower than what it could be theoretically.

Our question of interest is whether any of the empirically observed Manhattan distance values in the striatum is substantively greater than the rest, within the neurobiologically realized range of laterality. To examine this question, we tested for the presence of outliers in the voxelwise distribution of Manhattan distance values. To establish an outlier threshold we used the interquartile range (IQR) criterion[32], defined as values above the sum of the distribution’s third quartile and 1.5 multiplied by the interquartile range (i.e. > q_.75_ + (1.5*IQR)). These threshold values were determined to be Manhattan distance > 1.142 for the right striatum, and Manhattan distance > 1.167 for the left striatum. Voxels whose Manhattan distance was so classified as an outlier were labeled as “high laterality” (HL) voxels, and clusters of HL voxels were considered to constitute “laterality hotspots”.

#### Laterality Directionality

The Manhattan distance calculation, with its inclusion of absolute value terms, obscures whether overall connectivity laterality is stronger in the direction of the right or left frontal cortex. We therefore conducted an additional “directionality analysis” by repeating the above steps with an altered Manhattan distance calculation that excludes the absolute value computations. This additional voxel-wise map was used to determine the direction of laterality at hotspots identified in the initial Manhattan distance analysis.

#### Comparison of laterality in homotopic striatal areas

To compare the magnitude of laterality in homotopic areas of the right and left striatum in a voxel-wise manner, we carried out the following procedure. First, we used AFNI’s[33] 3dLRflip program on each subject’s right striatum Manhattan distance map, to align its voxels with the homotopic voxels (and therefore, the approximately homotopic structural areas) of the subject’s left striatum Manhattan distance map. We then took the difference of these two maps, masked by the intersection of the two masks to exclude voxels without a homotopic pair (19% of voxels in the union of the maps). This resulted in a final voxel-wise map for each subject whose values represent the Manhattan distance difference (i.e., difference in laterality) between homotopic striatal voxels. These subject-level maps were then averaged to create a group-level Manhattan distance difference map. Here, we again tested for the presence of HL voxels to identify voxels whose Manhattan distance difference was substantively different than the rest. The IQR threshold for this analysis was determined to be Manhattan distance difference > 0.464. To further corroborate findings, we conducted a paired t-test by comparing, in a voxel-wise manner, each subject’s right Manhattan distance heatmap to their left striatum Manhattan distance heatmap. This analysis examined the statistical significance of the average within-subject Manhattan distance difference at each homotopic voxel pair. We used a voxel-level threshold of p<0.001 with 3dClustSim (AFNI 20.1.14) to determine a p<0.05 family-wise corrected cluster-level threshold, corresponding to k>6.

#### Identification of frontal cortical ROIs driving laterality at hotspots

At each striatal hotspot’s peak Manhattan distance coordinates, we examined the absolute value of the difference between the Z-scores for each right hemisphere - left hemisphere ROI pair. These difference values measure the magnitude of laterality for each ROI pair. In order to determine the ROI pairs most strongly contributing to laterality at each hotspot, we identified ROI pairs whose |RH_Z_ – LH_Z_| value constituted an outlier in the distribution of all ROI |RH_Z_ – LH_Z_| values throughout the striatum. This threshold was |LH_Z_ – RH_Z_| > 0.076 for the right striatum and |LH_Z_ – RH_Z_| > 0.078 for the left striatum.

#### Replication Analysis

To examine replicability, we repeated the above analyses in a separate, age-, gender-, and handedness magnitude-matched sample of 77 HCP subject. We used nearest neighbor matching via the R program MatchIt (https://cran.r-project.org/web/packages/MatchIt/MatchIt.pdf) to select a replication sample matched on the aforementioned variables to the initial discovery sample. Briefly, the program computed a distance between each discovery sample subject and the remaining HCP subjects, and, one-by-one, selected a match for each discovery sample subject.

To evaluate the spatial correspondence between laterality hotspot voxels identified in the initial Discovery sample and those identified by the Replication sample, we performed a Dice coefficient analysis. To do so, we first calculated the Dice coefficient[34] between corresponding laterality hotspot voxels clusters, defined as

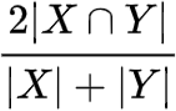

where |X ⋂ Y| is the number of voxels where cluster X and cluster Y overlap, |X| is the total number of voxels in cluster X, and |Y| is the total number of voxels in cluster Y. Then, to determine a significance threshold we performed permutation testing. Using the recently developed BrainSMASH[35] program to create permuted null brain maps that preserve spatial autocorrelation, on each of 10,000 iterations the voxel locations of cluster Y were randomized throughout the striatal mask, and a Dice coefficient between cluster X and the spatially randomized cluster Y were calculated. This created a Dice coefficient test statistic distribution customized to the striatal mask space. Clusters were considered to have significant spatial overlap if their Dice coefficient corresponded to p < 0.05.

#### Atlas Sensitivity Analysis

In order to ensure that laterality results were not an artifact of the specific atlas cortical parcellation schema chosen and were robust across parcellation definitions, we repeated the analyses using the Automated Anatomic Labeling (AAL) atlas (**Supplemental Figure 1b**), and again evaluated spatial correspondence using Dice coefficient analysis and permutation testing. If the connectivity profile laterality metric is indeed an inherent property of a voxel’s connectivity profile and not an artifact of a specific atlas parcellation, we would expect the voxels identified as having comparatively high laterality to remain largely consistent, regardless of the atlas used. This follows from how the Manhattan distance is calculated.

The Manhattan distance calculation involves a summing over all right-hemisphere/left-hemisphere ROI-pair differences in an atlas. The input data to the Manhattan distance calculation, therefore, is sampled from the full space of the atlas (i.e., data from all voxels in right frontal cortex and all voxels in left frontal cortex are included). If two atlases cover roughly the same underlying space (e.g., right and left frontal cortex), then using either atlas for a Manhattan distance calculation will include data from roughly the same space. This is because regardless of how the ROIs are parcellated on top of that space, the ROI-pair differences are summed back together at the end of the calculation. Thus, while a voxel’s Manhattan distance value itself would differ in different atlas parcellations, its relative magnitude compared to other voxels should remain consistent. For example, voxels with the highest Manhattan distance using one atlas parcellation should also have the highest Manhattan distance using another atlas parcellation.

#### Laterality in Right-handed versus Left-handed Individuals

Since all subjects in the discovery and replication samples were right-handed, we also repeated the analysis in an additional sample consisting only of left-handed individuals from the HCP. Similar laterality magnitude maps in right-handed and left-handed individuals would suggest that, regardless of which hemisphere is dominant, the magnitude of the hemispheric dominance in homotopic striatal areas is similar. Age, gender, and handedness information for each sample is presented in **Table 1**. Here, we performed a voxel-wise two-sample t-test to assess laterality magnitude differences between these left-handed individuals and the righthanded individuals from the discovery sample. Additionally, we performed supervised classification using a support vector machine (SVM) with MATLAB’s Statistics and Machine Learning Toolbox (version 2020b) to examine whether the voxel-wise Manhattan distance (i.e., laterality magnitude) maps could be used to distinguish between the right-handed and lefthanded individuals. To avoid data leakage and overfitting, we used nested 10-fold crossvalidation to estimate an unbiased generalization performance. First, the dataset was randomly divided into 10 folds of approximately 15 subjects each. Then, for each of 10 iterations, one of the 10 folds was left out for testing, while the other nine folds were used for model training. On each iteration, an outer loop performed dimensionality reduction via feature selection on the training set by selecting the 25% most predictive features (i.e., voxels), as determined via F-tests, for model training [36]. Then, for hyperparameter optimization and model selection, an inner loop performed Bayesian optimization with 10-fold cross validation on the training set using the selected features. Lastly, the optimized model was used to predict the handedness of the subjects in the left-out fold. The final metric of interest was the mean accuracy of test predictions across each of the 10 iterations.

**Table 1.** Sample Demographics

|  | Discovery Sample<br>(n=77) | Replication Sample<br>(n=77) | Left-handed Sample<br>(n=77) |
| --- | --- | --- | --- |
| Age (mean ± SD) | 29.09 ± 3.86 | 29.08 ± 3.62 | 28.75 ± 3.72 |
| Gender (% male) | 36.4% | 33.8% | 50.6% |
| Edinburgh Handedness<br>Inventory (mean ± SD) | 89.68 ± 8.79 | 90.00 ± 8.55 | -64.68 ± 21.34 |

#### Laterality and Behavior

To examine whether voxel-wise laterality magnitude was associated with performance on a language-related (i.e., lateralized) behavioral task, we conducted two types of analyses. First, we performed traditional, univariate voxel-wise regressions to assess whether the laterality magnitude of individual voxels was correlated with task performance. Second, we performed supervised support vector regression (SVR) to examine whether voxel-wise laterality magnitude maps could predict task performance. The model was trained on the Discovery sample, and then tested on the Replication sample. For training, we first selected the 25% most predictive voxels (determined via *F*-tests). We then used Bayesian optimization with ten-fold cross-validation to optimize hyperparameters. The optimized model trained to take the identified set of voxels as input was then used to predict the behavioral performance of subjects in the Replication sample. The outcome metric of interest was the correlation between actual and predicted performance.

## RESULTS

### Comparison of ipsilateral and contralateral frontal cortex fingerprints within the striatum

We identified several laterality “hotspots” indicative of comparatively high *dissimilarity* between ipsilateral and contralateral frontal cortical-striatal connectivity profiles. Laterality magnitude heatmaps (**Figure 1a**) illustrate peaks in the left rostral central caudate, the right rostral ventral putamen, and the right caudal ventral caudate. In the right striatum, a cluster of 13 HL voxels, constituting 0.68% of all right striatal voxels, was identified in the rostral ventral putamen (**Figure 1b**). At the peak Manhattan distance voxel of this cluster, the |RH_Z_ - LH_Z_| laterality of seven ROI pairs surpassed the outlier threshold (**Table 2**). Furthermore, the directionality analysis showed that this hotspot had the most right-lateralized frontal cortical connectivity of anywhere in the bilateral striatum (**Supplemental Figure 3**). One HL voxel was also identified in the caudal ventral caudate, where the |RH_Z_ - LH_Z_| laterality of nine ROI pairs surpassed the HL threshold (**Table 2**). In the left striatum, a cluster of 37 HL voxels, constituting 1.95% of all left striatal voxels, was identified in the rostral central caudate (**Figure 1b**). At the peak Manhattan distance voxel of this cluster, the |RH_Z_ - LH_Z_| laterality of eight ROI pairs surpassed the HL threshold (**Table 2**). Furthermore, the directionality analysis showed that this hotspot had the most left-lateralized frontal cortical connectivity of anywhere in the bilateral striatum (**Supplemental Figure 3**). The pars opercularis was the only ROI amongst the top three most lateralized ROIs at all three hotspots.

**Figure 1.**
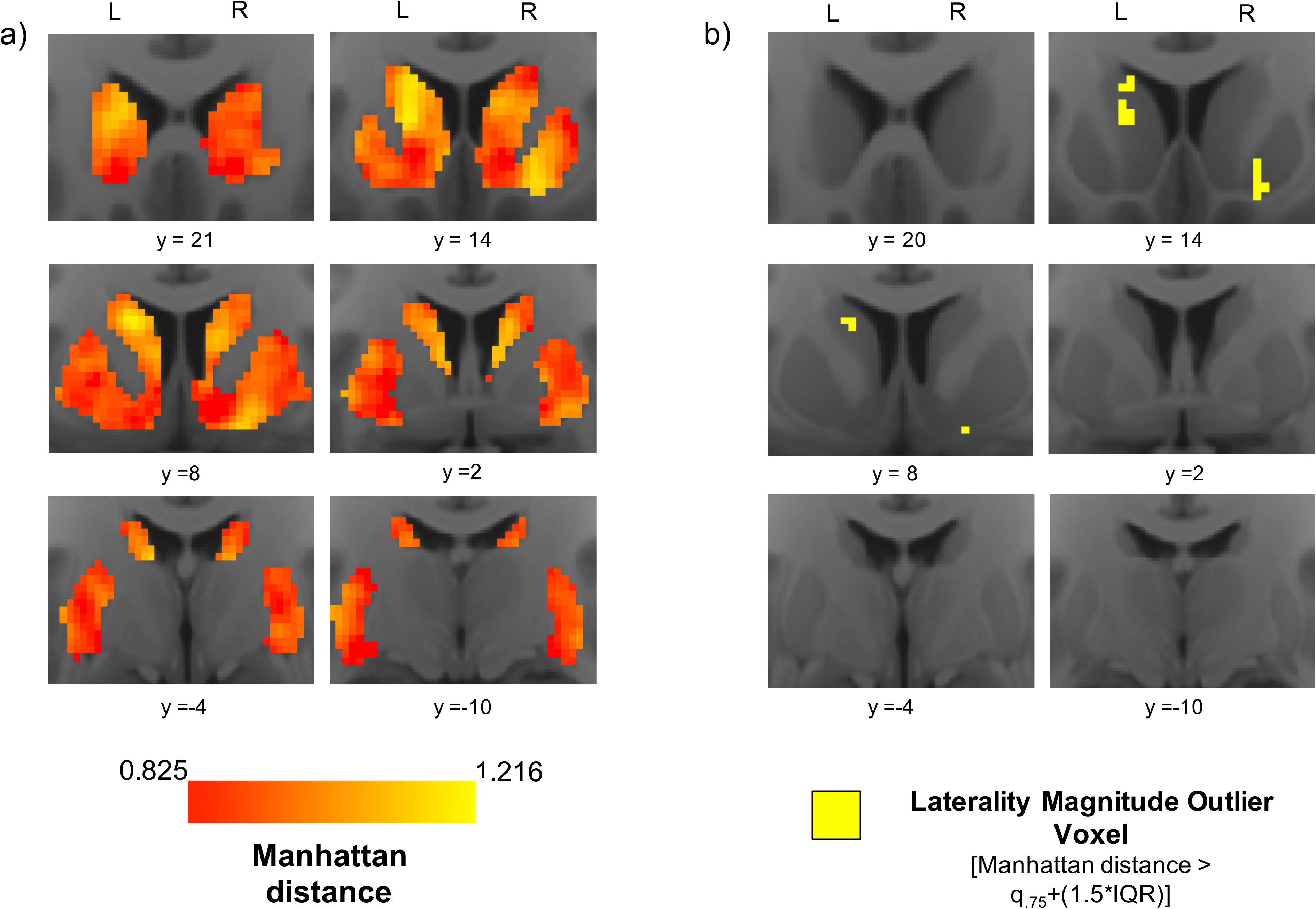
Functional connectivity profile laterality: heatmaps and hotspots. a) Average voxel-wise heatmaps of Manhattan distance in the left and right striatum from the Discovery data set. Warm colors indicate larger difference between a voxel’s ipsilateral and contralateral frontal cortical connectivity profile. b) Maps highlighting laterality “hotspot” voxels whose Manhattan distance constitutes an extreme (outlier) in the distribution.

**Table 2.** Frontal Cortical Drivers of Laterality

|  | Right Rostral<br>Ventral<br>Putamen<br>(16, 10, -14) | Left Rostral<br>Central<br>Caudate<br>(-12, 10, 16) | Right Caudal<br>Ventral<br>Caudate<br>(10, 4, 10) |
| --- | --- | --- | --- |
| Frontal Cortical ROI | Connectivity<br>Laterality<br>$\vert LH_Z - RH_Z\vert $ | Connectivity<br>Laterality<br>$\vert LH_Z - RH_Z\vert $ | Connectivity<br>Laterality<br>$\vert LH_Z - RH_Z\vert $ |
| Frontal Orbital | **0.174** | **0.106** | **0.077** |
| Subcallosal Cortex | **0.132** | **0.097** | **0.086** |
| Pars Opercularis | **0.105** | **0.134** | **0.109** |
| Pars Triangularis | **0.097** | **0.088** | **0.125** |
| Frontal Pole | **0.083** | **0.108** | **0.095** |
| Middle Frontal Gyrus | **0.078** | **0.122** | **0.090** |
| Superior Frontal Gyrus | **0.076** | **0.094** | **0.108** |
| Frontal Operculum | 0.074 | **0.089** | **0.078** |
| Frontal Medial | 0.064 | 0.063 | 0.057 |
| Insula | 0.056 | 0.056 | 0.056 |
| Paracingulate Gyrus | 0.053 | 0.070 | **0.077** |
| Supplementary Motor | 0.052 | 0.055 | 0.052 |
| Central Operculum | 0.045 | 0.043 | 0.049 |
| Precentral Gyrus | 0.043 | 0.048 | 0.041 |
| ACC | 0.038 | 0.043 | 0.043 |

Bolding indicates that an ROI's $|LH_Z - RH_Z|$ value at the hotspot peak coordinate surpassed the striatum-wide $|LH_Z - RH_Z|$ outlier threshold ( $|LH_Z - RH_Z| > 0.076$ for the right striatum and $|LH_Z - RH_Z| > 0.078$ for the left striatum).

### Pars Opercularis Laterality

Given observations of strong connectivity laterality with the pars opercularis at each of the striatal laterality hotspots, we mapped the voxel-wise laterality of pars opercularis connectivity throughout the striatum to visualize the complete spatial distribution of its magnitude (**Figure 2**). For this mapping, we used RH_Z_ – LH_Z_ rather than |RH_Z_ – LH_Z_| in order to distinguish between striatal areas with RH vs. LH pars opercularis-dominated laterality. We also examined ROI-specific HL voxels (IQR thresholds: RH_Z_ – LH_Z_ > ±0.076 for the right striatum and RH_Z_ – LH_Z_ > ±0.078 for the left striatum). In the right striatum, RH pars opercularis-dominated laterality peaked in the rostral ventral putamen; a cluster of 12 HL voxels was identified in this region, with no HL voxels identified elsewhere. There were no instances of LH pars opercularis-dominated HL voxels in the right striatum. In the left striatum, LH pars opercularis-dominated laterality peaked in the rostral central caudate; a cluster of 31 HL voxels was identified in this region, and no HL voxels were identified elsewhere. There were no instances of RH pars opercularis-dominated HL voxels in the left striatum.

**Figure 2.**
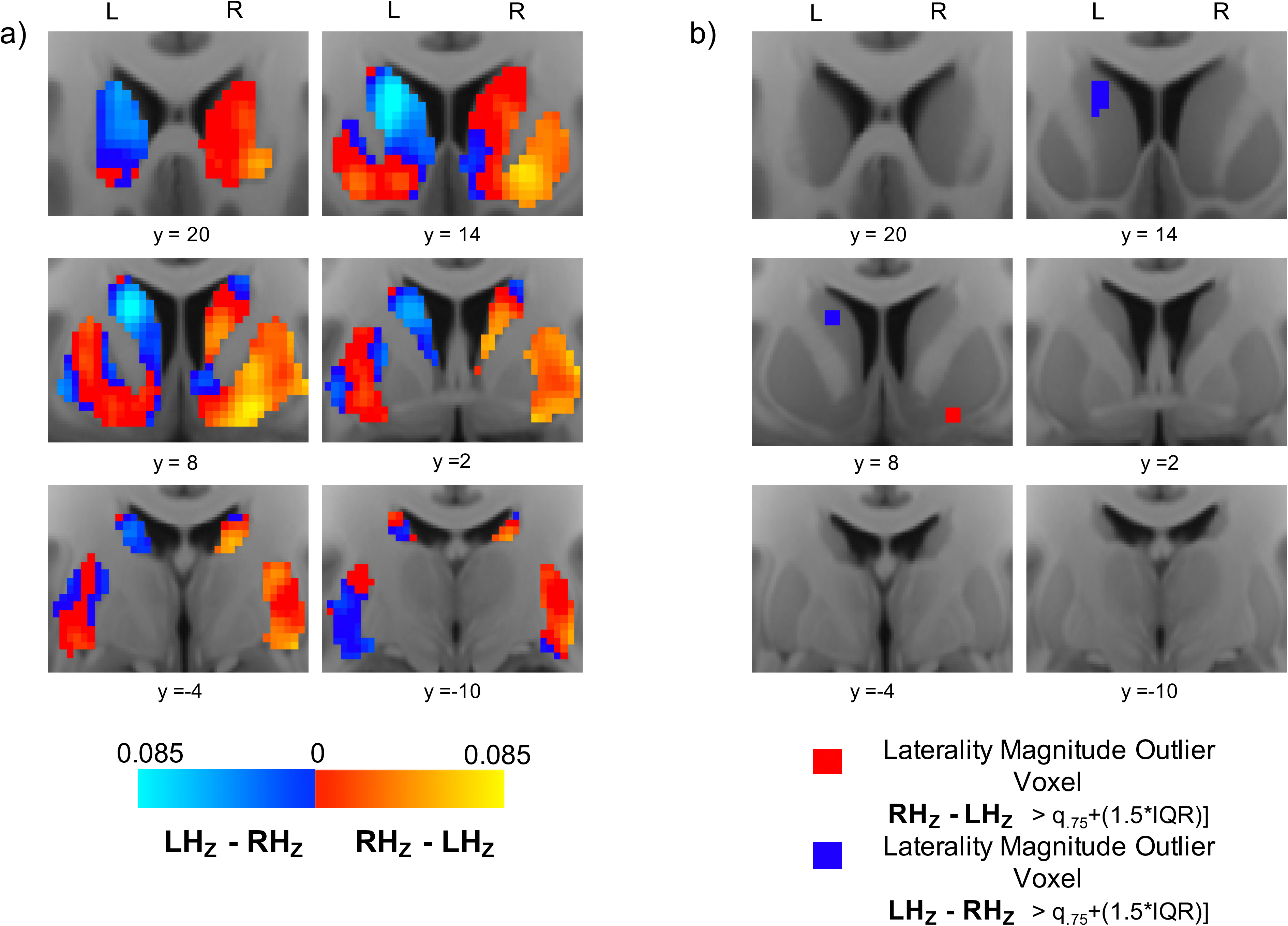
Pars opercularis connectivity laterality: heatmaps and hotspots. a) Average voxel-wise heatmaps of laterality in pars opercularis functional connectivity in the left and right striatum. Warm spectrum colors indicate voxels whose connectivity with right pars opercularis is stronger than their connectivity with left pars opercularis. Cooler spectrum colors indicate voxels whose connectivity with left pars opercularis is stronger than their connectivity with right pars opercularis. Brighter hues on each spectrum denote higher levels of laterality. b) Maps highlighting laterality “hotspot” voxels where the difference between connectivity with left and right pars opercularis constitutes an extreme (outlier) in the distribution

### Comparison of laterality magnitude in homotopic areas of the right and left striatum

The Manhattan distance difference heatmap illustrates peaks in the central caudate and ventral putamen (**Figure 3a**). A cluster of 5 HL voxels was identified in the central caudate, indicating that the difference in laterality magnitude between the left and right striatum is substantively larger in this region compared to the rest of the striatum (**Figure 3b**). Furthermore, the paired samples t-test revealed that the difference between left and right striatum laterality magnitude was statistically significant within a cluster of size k=11 with peak coordinates at (−14, 16, 14) in the central caudate, within a cluster of size k=16 with peak coordinates at (−18, 16, −10) in the rostral ventral putamen, and within a cluster of size k=10 with peak coordinates at (−26, 0, −8) in the caudal ventral putamen (**Figure 3c**).

**Figure 3.**
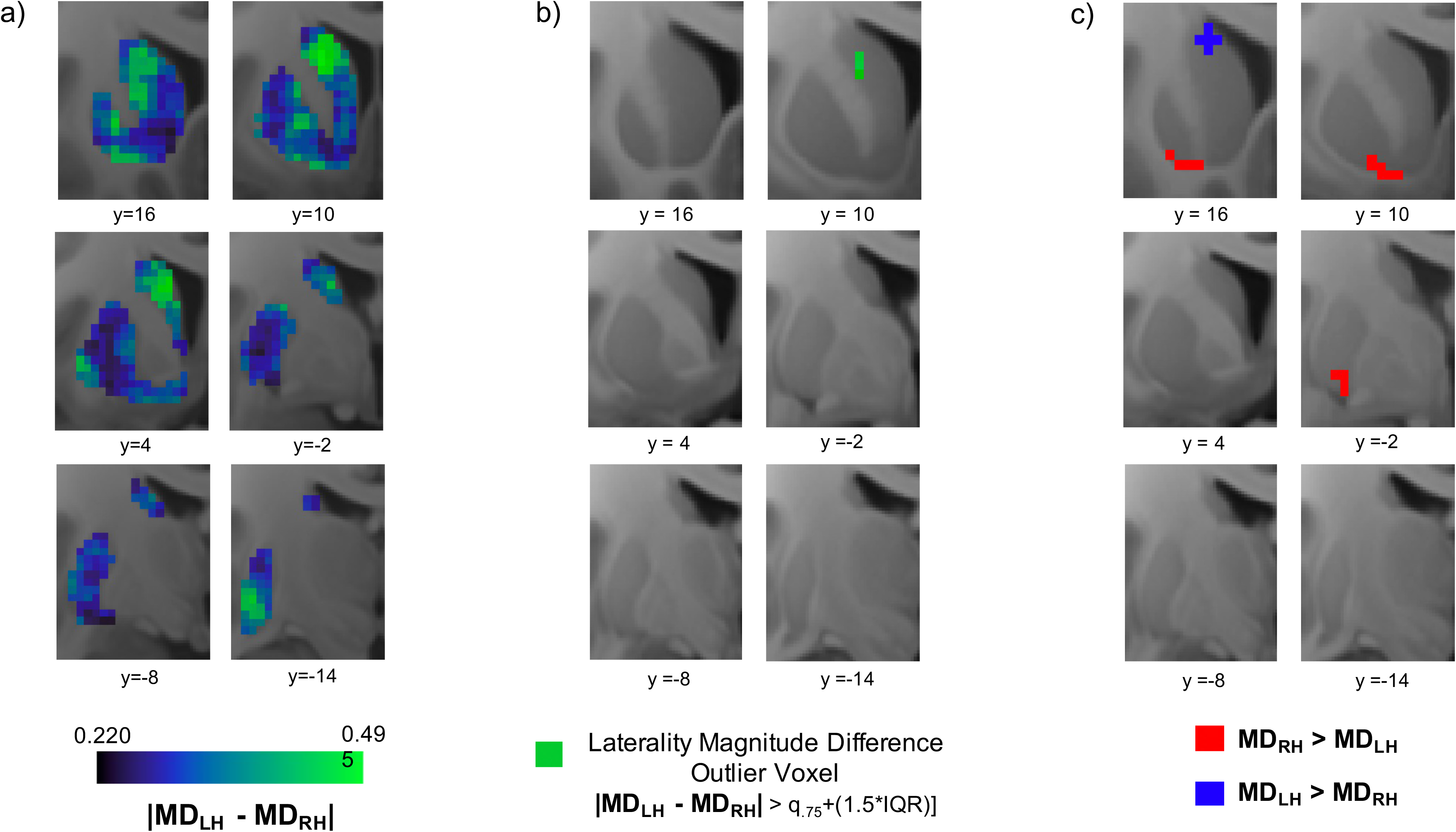
Laterality magnitude difference in homotopic striatal voxels. a) Voxel-wise heatmaps of Manhattan distance difference values. The value at each voxel represents the subject average difference in connectivity profile laterality between the voxel and its homotopic pair in the contralateral striatum. Greener colors indicate higher levels of Manhattan distance difference, and bluer colors indicate lower levels. b) Maps highlighting laterality difference “hotspot” voxels where the difference in laterality between the right and left striatum constitutes an extreme (outlier) in the distribution. c) Statistically significant clusters from paired t-test. Red clusters indicate areas where right striatum laterality magnitude was significantly larger than left striatum laterality magnitude; blue clusters indicate areas where left striatum laterality magnitude was significantly larger than right striatum laterality magnitude.

### Sensitivity and Replication Analyses

#### Replication Analysis

The identified HL voxels in the replication data had significant spatial correspondence to those identified in the Discovery sample right striatum (Dice coefficient = 0.095, p<0.0295) and left striatum (Dice coefficient = 0.329, p<0.0001), with overlap present in the right rostral ventral putamen and left rostral central caudate (**Figure 4a-b**).

**Figure 4.**
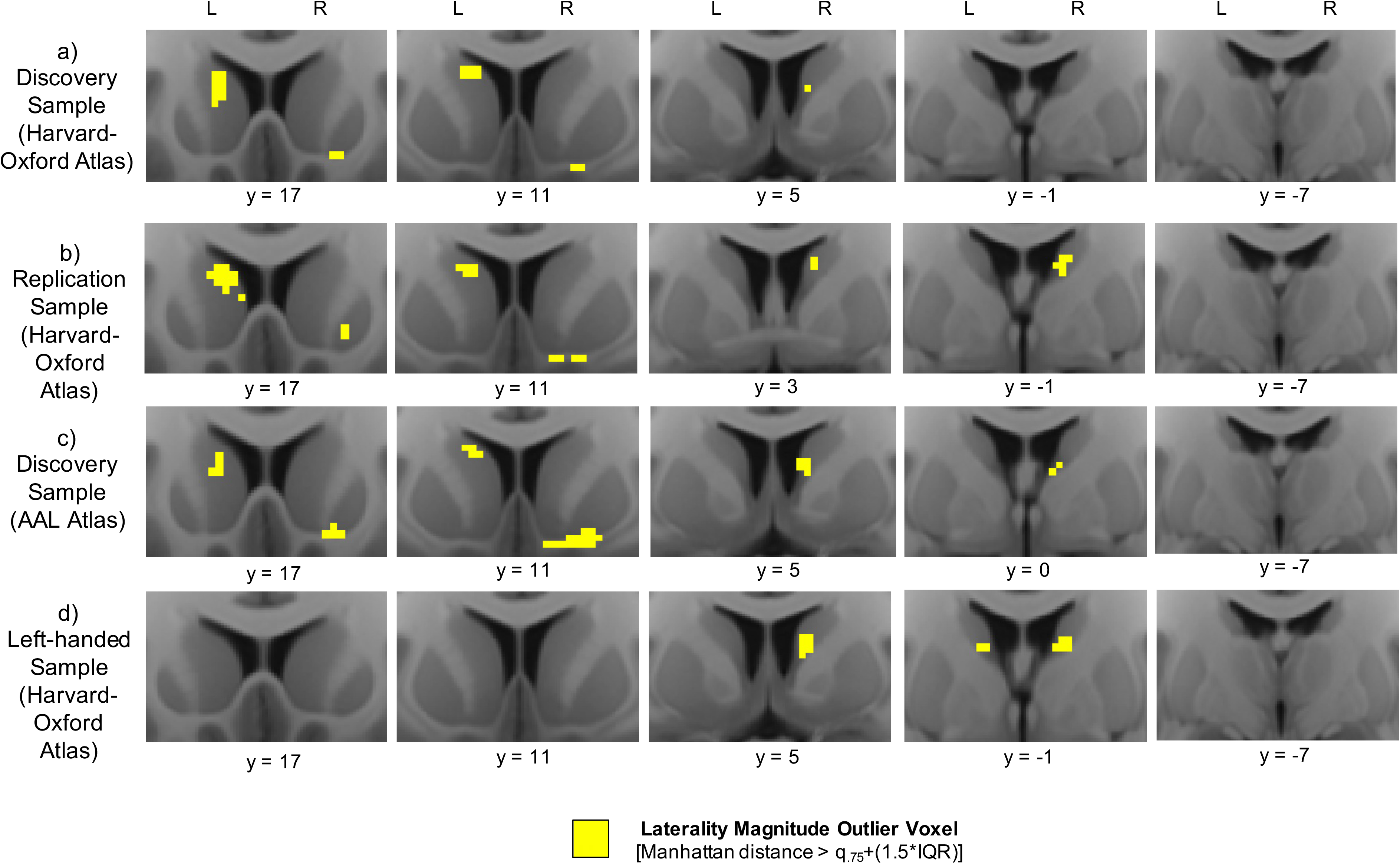
Replication and sensitivity analyses. Comparing laterality “hotspot” outlier voxels from the Discovery sample (a) to those from the Replication sample (b), the atlas sensitivity analysis (c), and the left-handed sample (d).

#### Atlas Sensitivity Analysis

In the atlas sensitivity analysis, identified HL voxels using the AAL atlas had significant spatial correspondence to those identified using the Harvard-Oxford atlas in both the right striatum (Dice coefficient = 0.333, p<0.0001) and left striatum (Dice coefficient = 0.607, p<0.0001), with overlap present in the right rostral ventral putamen and left rostral central caudate (**Figure 4a and 4c**).

#### Left-handed Replication Sample

As in the Discovery sample, HL voxels were identified in the left-handed cohort in the right caudal ventral caudate (**Figure 4d**). However, HL voxels were not identified in the right rostral ventral putamen or left rostral central caudate, and therefore spatial correspondence of HL voxels was not significant for either the right striatum (Dice coefficient = 0.041, p<0.206) or left striatum (Dice coefficient = 0, p = 1). Nonetheless, overall heatmaps were qualitatively similar (**Supplemental Figure 4**), the two-sample t-test did not reveal any significant voxel-wise group differences, and the machine learning classifier trained on the heatmaps was not able to distinguish between the two groups above chance level (53.9%).

#### Post-Hoc Analyses

The small size of the HL voxel clusters thresholded using the IQR outlier criterion led to small Dice coefficient values, despite qualitative evidence of high spatial similarity of corresponding clusters between groups (**Figure 4**). To further corroborate this similarity quantitatively, we first thresholded the clusters at less stringent values and re-calculated Dice coefficients (**Supplemental Table 1**) and constructed receiver-operating characteristic (ROC) curves by varying the threshold to assess area under the curve (AUC) (**Supplemental Figure 5**). The additional threshold values used were the maximum value, third quartile, median, first quartile, and minimum value of each group’s voxel-wise Manhattan distance distribution. Second, we assessed the correlation between each sample’s unthresholded Manhattan distance maps (**Supplemental Table 2**). High Dice coefficient values were observed for less stringent thresholds, and all unthresholded Manhattan distance maps were strongly and significantly correlated with one another.

### Laterality Magnitude and Behavior

Univariate voxel-wise analyses did not reveal any significant relationships between the laterality magnitude of individual voxels and the behavioral measures. However, the machine learning approach revealed that the multivariate heatmap of laterality magnitudes across voxels of the left striatum significantly predicted performance on the ReadEng task (r = 0.251, p = 0.028) (**Figure 5**). In contrast, right striatum laterality magnitude heatmaps were not significantly predictive of performance on the ReadEng task (r = 0.042, p = 0.718). Finally, laterality magnitude heatmaps were not significantly predictive of performance on the Delay Discounting task for either the right striatum (DDisc_AUC_200: r= −0.080, p=0.491; DDisc_AUC_40K: r= −0.032, p=0.784) or left striatum (DDisc_AUC_200: r < 0.001, p > 0.999; DDisc_AUC_40K: r = 0.096, p=0.408).

**Figure 5.**
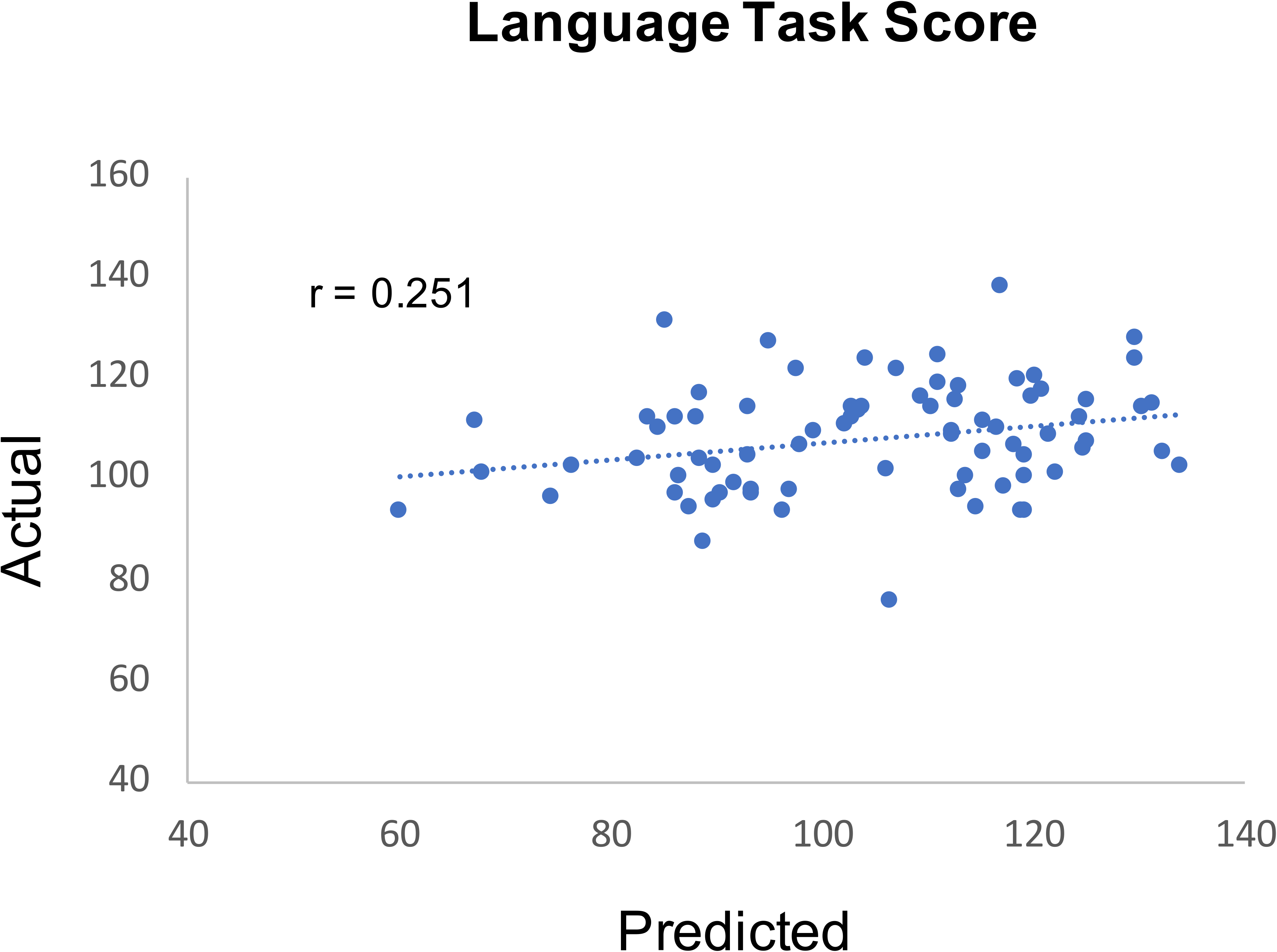
Laterality heatmap prediction of language task performance. The relationship between subjects’ actual performance on the ReadEng language task and predicted performance based on their left striatum voxel-wise laterality heatmap.

## DISCUSSION

Functional connectivity profile laterality has been shown to be very low throughout the human striatum [19, 20]. However, we identified a defined spatial distribution of laterality magnitude in which a handful of focal striatal areas display connectivity profiles that are substantively more lateralized than the rest of the striatum. These striatal “laterality hotspots” include the right rostral ventral putamen, the left rostral central caudate, and the right caudal ventral caudate. The robustness and spatial specificity of these hotspots was corroborated across several sensitivity and replication analyses. Furthermore, inter-individual variance in the striatum-wide distribution of connectivity profile laterality was predictive of performance on a task engaging language, a lateralized capacity, but not on Delay Discounting, a task engaging non-lateralized functional capacity.

Compared to the rest of the striatum, the identified striatal laterality hotspots appear to receive the most dissimilar information from the right and left frontal cortices, suggesting that they may play especially important roles in lateralized brain functions such as response inhibition and/or language. It is notable then that one of the largest sources of connectivity laterality at all of the laterality hotspots was the pars opercularis of the inferior frontal gyrus (BA44), itself a cortical region with well-established lateralized functionality as discussed below[37, 38].

In most right-handed (~95%) and left-handed (~75%) individuals[39, 40], BA44 in the left hemisphere, but not the right hemisphere, is part of Broca’s language area[41, 42]. The observation that BA44_LH_ - BA44_RH_ connectivity laterality peaks in the left central caudate suggests a particular role for this striatal area in language functions. Indeed, the left central caudate has been repeatedly implicated in several aspects of language including language acquisition[43], word-finding and the selection of appropriate lexical-semantic responses[44], and monitoring and controlling the language currently in use in bilingual speakers[45]. The comparatively lower BA44_LH_ - BA44_RH_ connectivity laterality in the homotopic right central caudate may reflect the more dominant role of the left central caudate in language processes.

On the other hand, BA44 in the right hemisphere is strongly implicated in motor inhibition processes[3, 4]. The observation that BA44_RH_ – BA44_LH_ connectivity laterality peaks in the right ventrolateral putamen may reflect a particular role for this striatal area in motor inhibition. This is consistent with a number of studies that find concurrent activation of right BA44 and right ventrolateral putamen during performance of response inhibition tasks[3, 46–50]. However, these studies also show involvement of the left ventrolateral putamen, which we find here also has stronger functional connectivity with right BA44 than left BA44. Overall, these findings suggest bilateral involvement of ventrolateral putamen in action inhibition might be related to outsized functional connectivity with right BA44.

Interestingly, connectivity profile laterality magnitude did not differ significantly between left-handed and right-handed individuals for any striatal voxels, and the machine learning classifier was not able to use striatum-wide laterality heatmaps to distinguish handedness above chance level. As in the right-handed groups, the left-handed group displayed HL voxels in the right caudal ventral caudate hotspot. And while voxels in the right rostral ventral putamen and left rostral central caudate did not surpass the HL threshold in the lefthanded group, voxels in these areas displayed comparable levels of laterality to those in the right-handed groups (**Supplemental Figure 4**). Furthermore, laterality maps as a whole for the right-handed and left-handed groups were strongly correlated (**Supplemental Table 1**). Overall, these findings suggest that, regardless of differences in intrinsic hemispheric dominance, the degree to which ipsilateral-contralateral frontal cortical connectivity differs at corresponding striatal loci is similar in the general population regardless of handedness. In tandem with the sensitivity analysis demonstrating that findings remain consistent across cortical atlas parcellations, these findings provide a robust, normative basis for future comparison with connectivity profile laterality in clinical samples.

The absence of univariate relationships between language task performance and connectivity profile laterality magnitude in focal striatal regions, despite the ability of a machine learning classifier to significantly predict performance based on striatum-wide laterality maps, suggests a complex and distributed brain-behavior relationship. Still, the fact that the classifier could only significantly predict language task performance from the left striatum laterality maps, and not from the right striatum laterality maps, is consistent with the predominantly left hemispheric lateralization of language function[39, 40].

In sum, we find that meaningful variation in functional connectivity profile laterality – both spatially within the striatum and across subjects – is evident in corticostriatal circuits. The elucidation of striatal laterality “hotspots” and their frontal cortical drivers warrants further examination of these sites’ roles in normative behavior and potential aberrance in psychiatric illness.

## Acknowledgements

Funded by National Institute on Drug Abuse grant 1F32DA048580-01A1 to CK and the Intramural Research Program of the National Institute on Drug Abuse (TR and EAS). Data were provided by the Human Connectome Project, WU-Minn Consortium (Principal Investigators: David Van Essen and Kamil Ugurbil; 1U54MH091657) funded by the 16 NIH Institutes and Centers that support the NIH Blueprint for Neuroscience Research; and by the McDonnell Center for Systems Neuroscience at Washington University.

