## Supplemental Materials for "Connectivity profile laterality in corticostriatal functional circuitry: a fingerprinting approach"

*Table S1. Dice coefficients between sample Manhattan distance maps at varying thresholds*

a)

| **Right Striatum** | Discovery and Replication | Discovery  and  AAL | Discovery and  Left-Handed |
| --- | --- | --- | --- |
| IQR Outlier Threshold | 0.095 | 0.333 | 0.041 |
| Quartile 3 Threshold | 0.664 | 0.812 | 0.530 |
| Median Threshold | 0.706 | 0.857 | 0.676 |
| Quartile 1 Threshold | 0.818 | 0.897 | 0.813 |

b)

| **Left**  **Striatum** | Discovery and Replication | Discovery  and  AAL | Discovery and  Left-Handed |
| --- | --- | --- | --- |
| IQR Outlier Threshold | 0.329 | 0.607 | 0 |
| Quartile 3 Threshold | 0.702 | 0.823 | 0.553 |
| Median Threshold | 0.763 | 0.878 | 0.679 |
| Quartile 1 Threshold | 0.818 | 0.939 | 0.796 |

*Table S2. Correlation between sample Manhattan distance maps*

a)

| **Right Striatum** | Discovery | Replication | AAL | Left-Handed |
| --- | --- | --- | --- | --- |
| Discovery | - | R=0.68, p<0.0001 | R=0.87,  p<0.0001 | R=0.62,  p<0.0001 |
| Replication | R=0.68, p<0.0001 | - | R=0.65,  p<0.0001 | R=0.61,  p<0.0001 |
| AAL | R=0.87,  p<0.0001 | R=0.65,  p<0.0001 | - | R=0.57,  p<0.0001 |
| Left-Handed | R=0.62,  p<0.0001 | R=0.61,  p<0.0001 | R=0.57,  p<0.0001 | - |

b)

| **Left**  **Striatum** | Discovery | Replication | AAL | Left-Handed |
| --- | --- | --- | --- | --- |
| Discovery | - | R=0.76,  p<0.0001 | R=0.93,  p<0.0001 | R=0.60,  p<0.0001 |
| Replication | R=0.76,  p<0.0001 | - | R=0.75,  p<0.0001 | R=0.64,  p<0.0001 |
| AAL | R=0.93,  p<0.0001 | R=0.75,  p<0.0001 | - | R=0.55,  p<0.0001 |
| Left-Handed | R=0.60,  p<0.0001 | R=0.64,  p<0.0001 | R=0.55,  p<0.0001 | - |

**Supplemental Figures Legend**

Figure S1. Frontal cortex parcellations for (a) the Harvard-Oxford atlas and (b) the Automated Anatomical Labeling (AAL) atlas. Colors are randomized and do not represent homologies or correspondences between atlases.

Figure S2. Analytical pipeline for calculating subject-level Manhattan distance maps.

Figure S3. Results of laterality directionality analysis, illustrating which voxels have stronger connectivity with the right frontal cortex (warm colors) or the with the left frontal cortex (cool colors).

Figure S4. Unthresholded group-level Manhattan distance maps for (a) the left-handed replication sample and (b) the right-handed discovery sample.

Figure S5. Dice coefficients between sample Manhattan distance maps at varying thresholds.
