## Supplemental Figures for "Connectivity profile laterality in corticostriatal functional circuitry: a fingerprinting approach"

### Slide 1
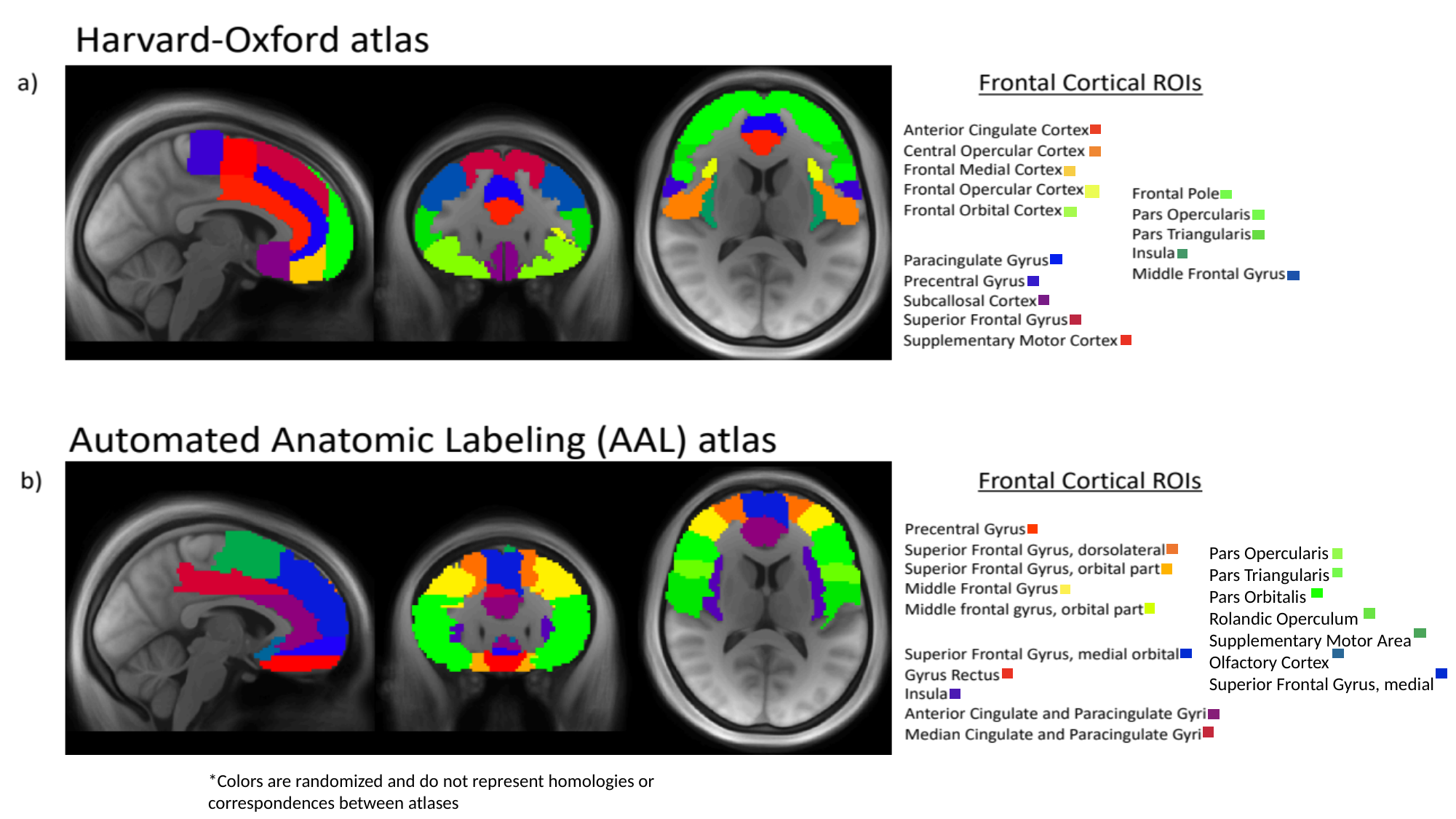

Pars Opercularis
Pars Triangularis
Pars Orbitalis
Rolandic Operculum
Supplementary Motor Area
Olfactory Cortex
Superior Frontal Gyrus, medial
*Colors are randomized and do not represent homologies or correspondences between atlases

### Slide 2
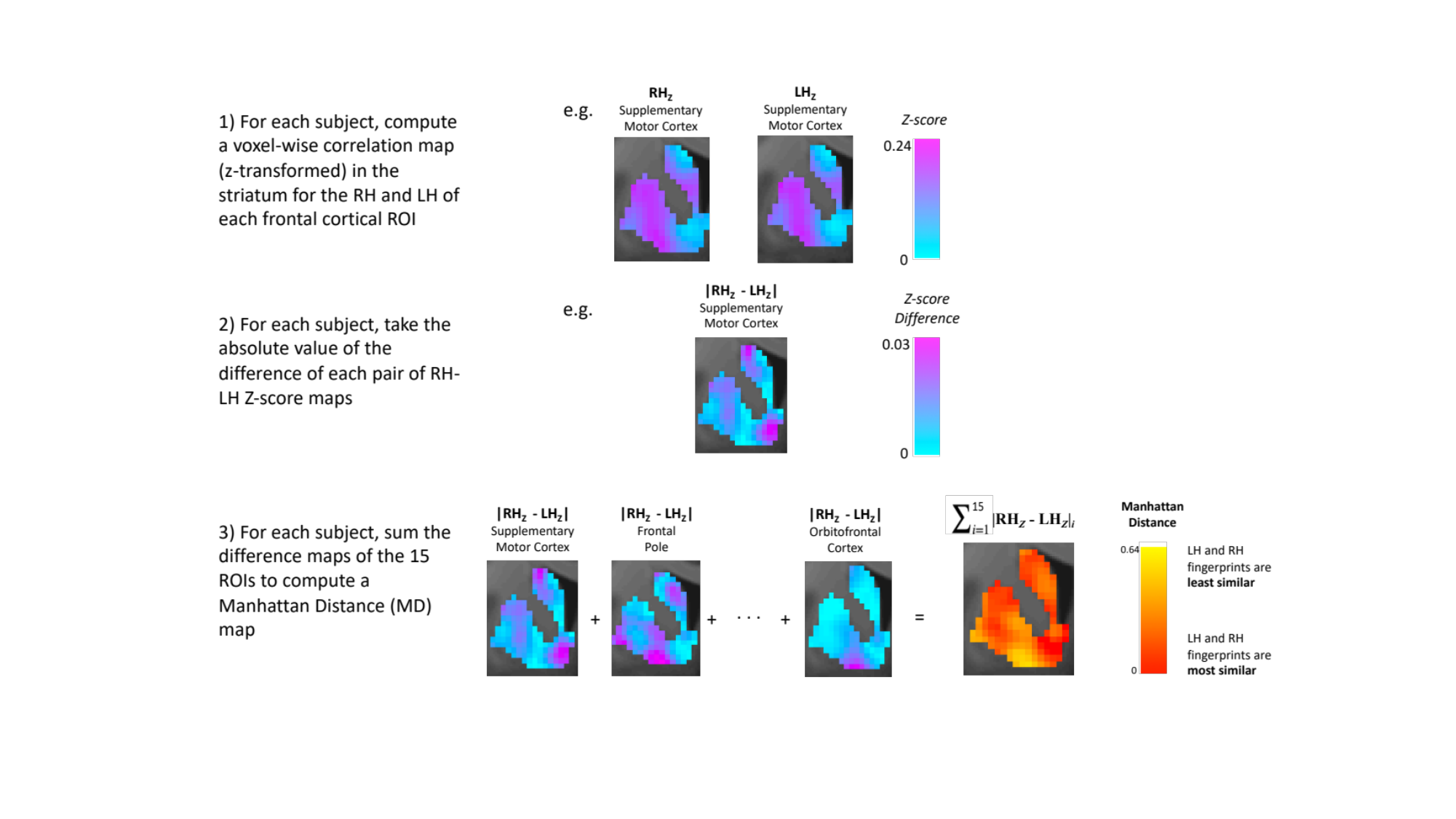

### Slide 3
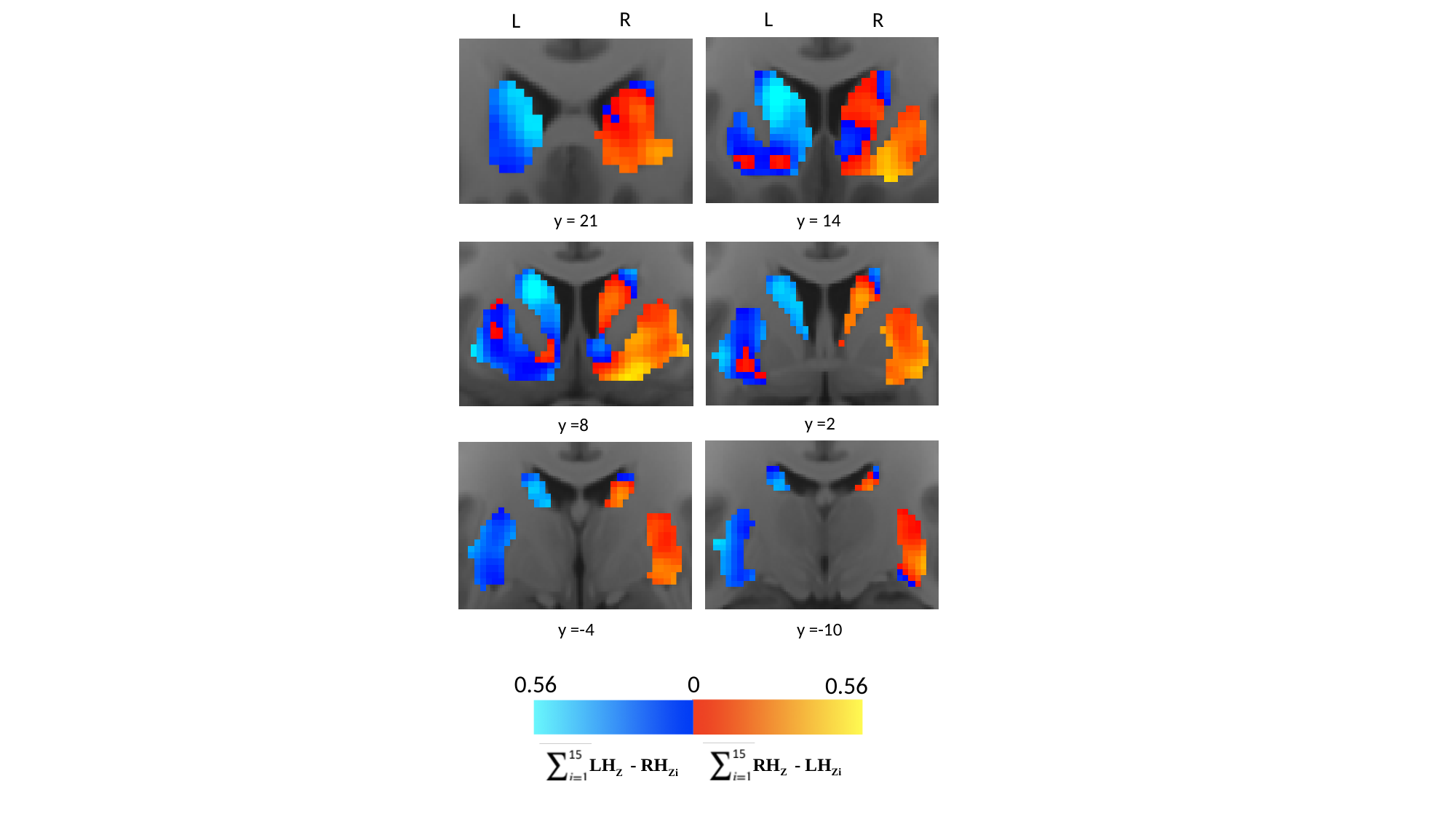

R
L
R
L
y = 14
y = 21
y =2
y =8
y =-10
y =-4
0.56
0
0.56
RH_Z_ - LH_Zi_
LH_Z_ - RH_Zi_

### Slide 4
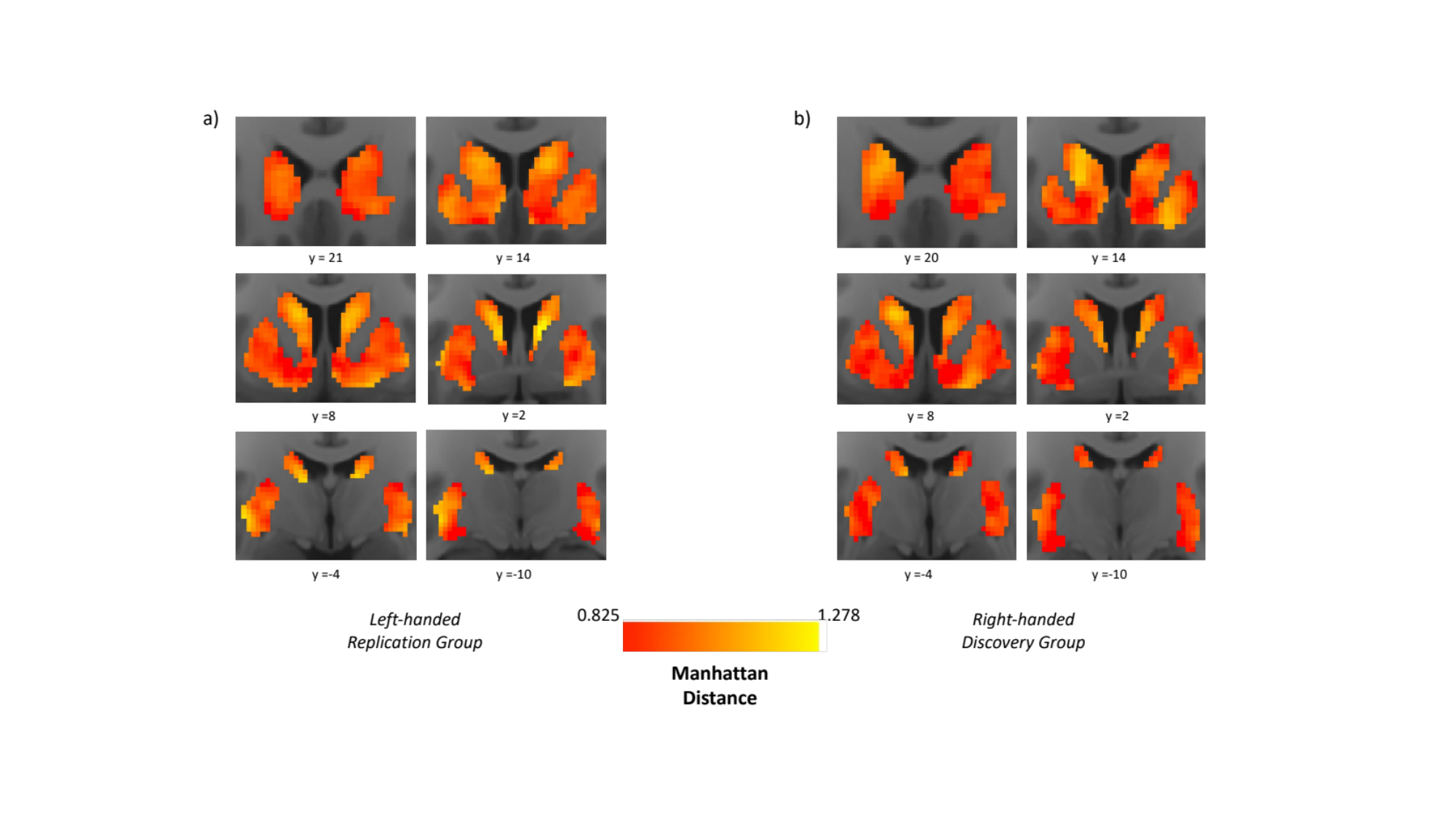

### Slide 5
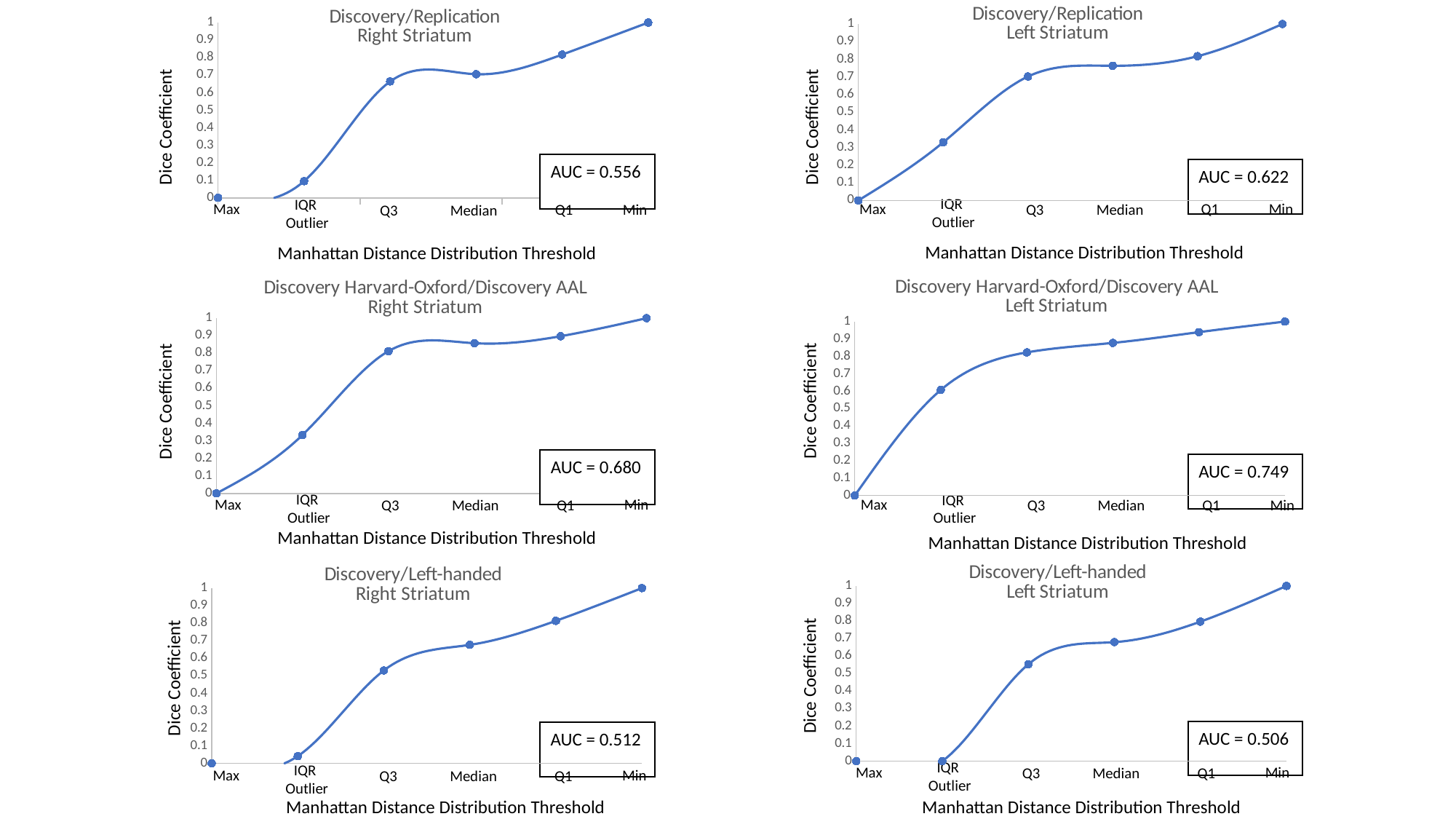

#### Chart: Discovery/Replication
Left Striatum
| Category |
|---|
#### Chart: Discovery/Replication
Right Striatum
| Category | |
|---|---|Dice Coefficient
Dice Coefficient
AUC = 0.556
AUC = 0.622
IQR
Outlier
Max
Min
Q1
Q3
Median
IQR
Outlier
Max
Min
Q1
Q3
Median
Manhattan Distance Distribution Threshold
Manhattan Distance Distribution Threshold
#### Chart: Discovery Harvard-Oxford/Discovery AAL
Right Striatum
| Category |
|---|
#### Chart: Discovery Harvard-Oxford/Discovery AAL
Left Striatum
| Category | |
|---|---|Dice Coefficient
Dice Coefficient
AUC = 0.680
AUC = 0.749
IQR
Outlier
Max
Min
Q1
Q3
Median
IQR
Outlier
Max
Min
Q1
Q3
Median
#### Chart: Discovery/Left-handed
Left Striatum
| Category |
|---|
#### Chart: Discovery/Left-handed
Right Striatum
| Category | |
|---|---|Manhattan Distance Distribution Threshold
Manhattan Distance Distribution Threshold
Dice Coefficient
Dice Coefficient
AUC = 0.506
AUC = 0.512
IQR
Outlier
Max
Min
Q1
Q3
Median
IQR
Outlier
Max
Min
Q1
Q3
Median
Manhattan Distance Distribution Threshold
Manhattan Distance Distribution Threshold
